# Microbiota-dependent and independent production of L-dopa in the gut of *Daphnia magna*

**DOI:** 10.1101/2021.08.02.454848

**Authors:** Rehab El-Shehawy, Sandra Luecke-Johansson, Anton Ribbenstedt, Elena Gorokhova

## Abstract

The host-microbiome interactions are essential for the physiological and ecological performance of the host, yet these interactions are challenging to identify. Neurotransmitters are commonly implicated in these interactions, but we know very little about the mechanisms of their involvement, especially in invertebrates. Here, we report a peripheral Catecholamine (CA) pathway involving the gut microbiome of the model species *Daphnia magna*. We demonstrate that: (1) tyrosine hydroxylase and dopa decarboxylase enzymes are present in the gut wall; (2) DOPA decarboxylase gene is expressed in the gut by the host, and its expression follows the molt cycle peaking after ecdysis; (3) biologically active L-Dopa, but not Dopamine, is present in the gut lumen; and (4) gut bacteria produce L-Dopa in a concentration-dependent manner when provided L-Tyrosine as a substrate. Impinging on gut bacteria involvement in host physiology and ecologically relevant traits, we suggest L-Dopa as a communication agent in the host-microbiome interactions in daphnids and, possibly, other crustaceans.

## Introduction

Host-microbiome interactions, including the role of gut bacteria in regulating host homeostasis and feedbacks between the microbiome and its host, are a hot topic in current evolutionary, ecological, and biomedical studies. Inter-kingdom communication is the bi-directional flow of signals between the host neurophysiological system and its microbiome, with neurotransmitters acting as signaling molecules in the host-microbiome metabolic axis. Bacteria recognize and produce common vertebrate neurotransmitters [1–4], such as catecholamines (CA) – L-Dopa, Dopamine, Epinephrine, and Norepinephrine [4]. L-Dopa (l-3,4 dihydroxyphenylalanine) is the precursor for Dopamine. Although L-Dopa itself is a neurotransmitter/modulator with receptors in the central and peripheral nervous system [5], its role as a signaling molecule *per se* in inter-kingdom communication is unclear. Mutualistic bacteria have been found responding to L-Dopa variation, e.g., Enterobacteriaceae and Pseudomonadaceae increase their growth *in-vitro* when supplemented with L-Dopa [6]. Moreover, several bacterial taxa produce L-Dopa *in-vivo*, and the microbial enzymes tyrosinase, tyrosine phenyl lyase, and *p*-hydroxyphenylacetate 3-hydroxylase are exploited in the biotechnological L-Dopa production [7]. Thus, bacteria recognize, respond and produce L-Dopa.

Animal guts are rich in CA and harbor commensal bacteria, especially in the epithelium-associated biofilms. Therefore, active CA-mediated communication between the gut microbiota and the animal host is likely to occur [4]. Of particular challenge is understanding molecular mechanisms and pathways of the host-microbiome interactions in organisms of varying complexity, moving away from mammalian studies to other relevant model species in ecology and evolution research. Although our knowledge of microbiome-mediated regulation of host development, immunity, homeostasis, and behavior progresses beyond the established model animals, most of what we know about host-microbiome interactions is based on vertebrate models, mainly mammals whereas studies with invertebrates are limited. Among the invertebrate models are branchiopod crustaceans in the genus *Daphnia* that are broadly used in ecology and evolution studies, including host-microbiome interactions [8–11]. *Daphnia* microbiome modulates the life cycle, including growth, reproduction, and tolerance to environmental stressors [8, 10, 12, 13], yet the molecular basis of the interactions affecting these ecologically and evolutionary relevant traits is poorly understood. Here, we hypothesized that CA are involved in host-microbiome interactions in *Daphnia magna*, and these interactions occur at the gut-lumen interface (Fig. 1). The hypothesis was tested using a series of experiments with *D. magna* to detect L-Dopa and Dopamine in the gut lumen, localize the enzymes catalyzing the first steps of the CA-pathway, follow the decarboxylation of L-Dopa to Dopamine by the host during the molt cycle, and evaluate the gut microbiota ability to produce L-Dopa. Collectively, these data were used to understand L-Dopa production in the gut by the host and its microbiome.

**Figure 1.**
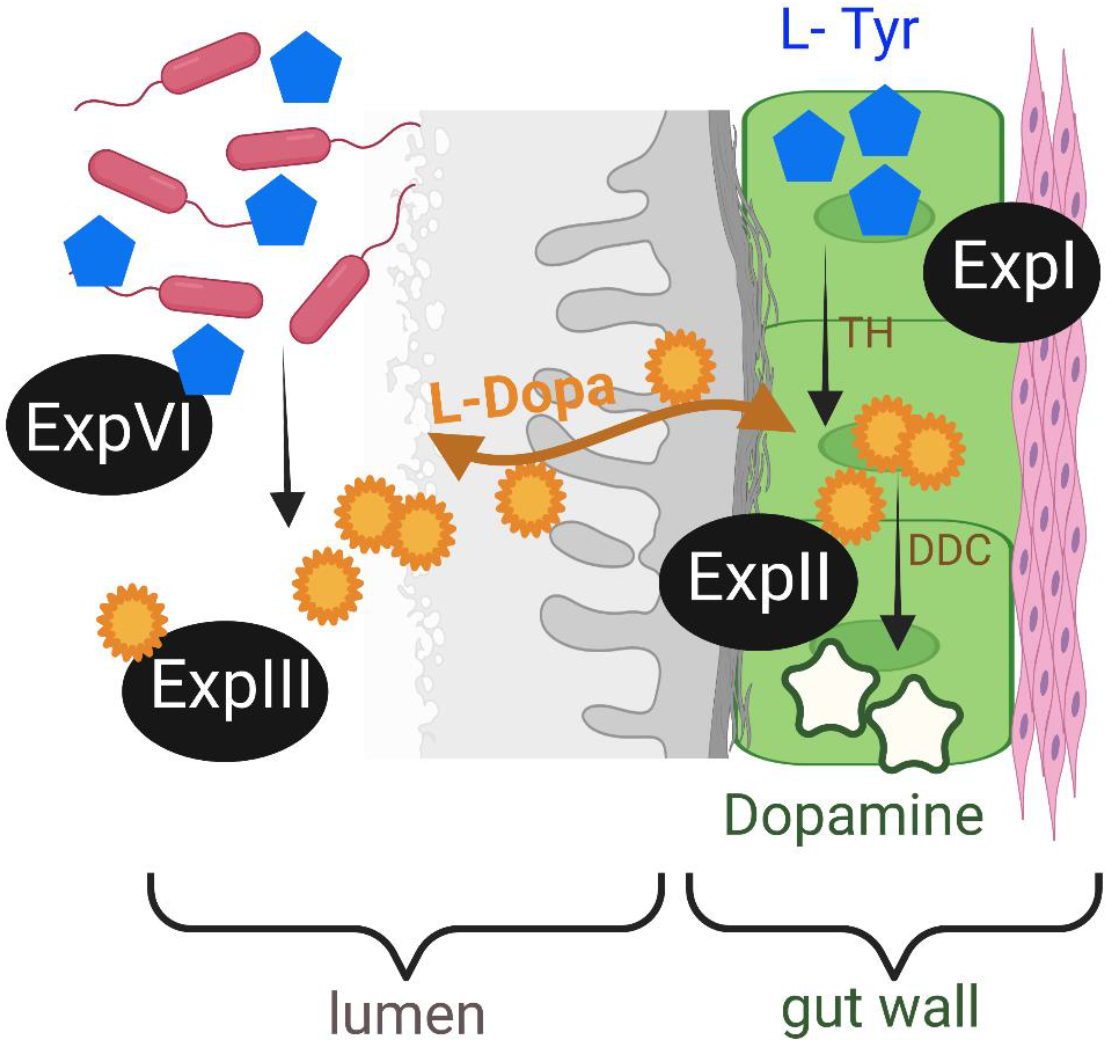
Conceptual diagram presenting the hypothesis and the associated experiments. The first step in the eukaryotic CA pathway is the hydroxylation of L-Tyrosine (L-Tyr) to L-Dopa via Tyrosine hydroxylase (TH). It is the rate-limiting step followed by the decarboxylation of L-Dopa to Dopamine by Dopa decarboxylase (DDC). L-Dopa is of dual origin and is an info-signal in a putative bi-directional communication between *Daphnia* and its gut bacteria in the lumen.

## Materials and methods

### Experimental design

The experimental design focused on the identification of the key reactions in the CA-pathway, i.e., the hydroxylation of L-Tyrosine to L-Dopa and the decarboxylation of L-Dopa to Dopamine, and the evaluation of the possible contribution of L-Dopa of bacterial origin to the total pool of L-Dopa in the *Daphnia* gut. Altogether, four experiments (Exp I-IV; Fig. 1) were conducted, with the purpose of:

– **Exp I**: localization of the CA-pathway in the gut wall using immunohistochemistry to confirm hydroxylation of L-Tyrosine, which is responsible for hydroxylation of L-Tyrosine to L-Dopa, and dopa decarboxylase (DDC) carrying out the decarboxylation of L-Dopa to Dopamine;
– **Exp II:** obtaining time-series of the expression of *Ddc* gene in the gut over the molt cycle using qPCR assay;
– **Exp III:** L-Dopa and Dopamine detection in *Daphnia* lumen using LC-HRMS;
– **Exp IV**: evaluation of the L-Dopa production by gut microbiota using enriched cultures of gut bacteria supplemented with L-Tyrosine to stimulate the growth of taxa capable of utilizing this substrate and measuring L-Dopa with LC-HRMS.

### *Daphnia magna* culture and standardization of test animals

All experiments were conducted with a single clone of *Daphnia magna* Straus (Cladocera, Branchiopoda; clone V, obtained from the Federal Environment Agency, Berlin, Germany). The animals are cultured under standard conditions [14] in M7 medium in groups of 12 ind. L^-1^ at 20 ± 2°C and 16:8 h light:dark cycle. The food, a mixture of green algae, *Pseudokirchneriella subcapitata* and *Scenedesmus spicatus*, was provided three times a week, and the medium was changed once a week.

Under these conditions, the molt cycle duration in Instars III to VI (the developmental stages used in this study) was 1.8 to 3.2 days. The animals used for immunostaining in Exp I were standardized with regard to their molt and embryo development stages: only Instars V-VI in the premolt stage carrying black-eyed embryos were used. The molt stage chronology was established in pilot experiments and found to follow well the published schedules [15, 16]. In Exp II, this chronology was used to assign the test animals to three main stages – postmolt, intermolt, and premolt, with midpoints 28%, 60%, and 85% of the molt cycle duration, respectively.

In daphnids, the digestive tract consists of an ectodermal foregut, an endodermal midgut, and an ectodermal hindgut. Prior to the gut sample collections for all four experiments, the daphnids were allowed to swim in sterile M7 for 10 min as a washing step and transferred to a sterile microscopy slide. The gut tube without the caeca (hereafter referred to as a gut sample) was dissected in each individual using sterile forceps and dissection needles.

### Localization of TH and DDC in the gut wall

In Exp I, immunohistochemistry was used to localize tyrosine hydroxylase (TH) in the daphnid gut wholemount; only premolt animals were used for this analysis (n = 7). As primary antibodies, we used Rabbit-polyclonal antibodies against TH (50997, Nordic BioSite; previously used to identify dopaminergic neurons in *D. magna* [17]) and Goat-polyclonal antibodies against Dopa decarboxylase (DDC) (AF3564, Novus Biologicals). Cross-absorbed Alexa-Fluor^488^ goat anti-rabbit IG (H+L) (A32731, ThermoFisher Scientific) and cross-absorbed Alexa-Fluor^488^ donkey anti-goat IgG (H+L) (A11055, ThermoFisher Scientific) were used as secondary antibodies. Negative controls without the primary antibodies or using IgG isotypes (Goat IgG NB410-28088, and Rabbit IgG NBP2-24891, Novus Biologicals) at the same dilution as the primary antibodies were used.

Immunolabelling was carried following the established methods [18, 19] with some modifications. In brief, the fixation of the dissected guts was carried out overnight in 4% paraformaldehyde. Samples were incubated with 5% BSA overnight (Super blocker, ThermoFisher Scientific) before applying primary antibodies. Incubation with primary antibodies was conducted for 48 h in the presence of 1% BSA (Super blocker, ThermoFisher Scientific). Incubation with secondary antibodies was conducted for 24 h in the presence of 1% BSA. The concentrations of antibodies were 1:2000 for both primary and secondary antibodies. The samples were rinsed 3 × 10 min, followed by 3 × 1 h, then 1 × overnight using PBS-TX (0.5%). All incubation and washing steps were done at 4°C on an orbital shaker in the dark.

Single images and Z-scans were taken using Zeiss Confocal Microscope 710 equipped with Aragon laser and 20× and 40× magnification and the manufacturer software. Images were processed using ImageJ2 [20]; no image manipulation was applied.

### RNA extraction, RT-qPCR, and gene expression analysis

In Exp II, synchronized cohorts were used to generate samples for premolt, intermolt, and postmolt stages. Total RNA was extracted from the gut samples (7-10 guts/sample) using the RNeasy Mini kit (Qiagen) and the on-column DNase I treatment (79254, Qiagen) according to the manufacturer instructions with additional in-tube DNase I treatment (AMPD1, Sigma-Aldrich). We used G3PDH (glyceraldehyde 3-phosphate dehydrogenase) as a housekeeping gene, which has a stable expression in *Daphnia* [21]. The primers for *Ddc* and *G3PDH* assays were adopted from Campos *et al*. [22] (Table S1); both primers were used to control for any residual DNA contamination. The RNA was reverse-transcribed using the High-Capacity RNA-to-cDNA™ Kit (ThermoFisher Scientific) following the manufacturer instructions. The qPCR assays were performed using QuantiNova SYBR green (Qiagen) as follows: cDNA (1µl, equivalent to 5 ng), forward and reverse primers (1 µM), 2 × QuantiNova SYBR Green PCR Master Mix (5 µl), QuantiNova ROX Reference Dye (1 µl), and DNA/RNA-free water (2 µl; Sigma-Aldrich) at 95°C for 2 min, 35 cycles at 95°C for 20 sec, and 60°C for 20 sec for annealing and data acquisition. A melt curve was generated after each run to ensure the reaction specificity. All qPCR runs were performed using Applied Biosystems™ StepOne™ Real-Time PCR system. The ΔCt values were calculated [23] to estimate the relative *Ddc* gene expression.

### *In-vitro* L-Dopa synthesis by *Daphnia* gut microbiome

In Exp IV, pre-enrichment was carried out by incubating whole guts in LB for 48 h at 24 °C with shaking. The overnight LB-cultures were used to inoculate (10% inoculum, vol/vol) test mixtures containing L-Tyrosine at concentrations 0, 1, 2 and 3 mM in M7; each concentration treatment was in triplicate. The inoculated cultures were then incubated for 24 h at 24°C with shaking. At the end of the incubation time, the culture density was measured at OD600nm. Sample preparation for LC-HRMS was carried as follows: 100 µl culture were transferred into a tube containing 300 µl of an ice-cold solvent mix of acetonitrile (ACN) and MilliQ mixtures, 85:15, with addition of 0.1% (vol/vol) formic acid (FA), followed by quick vortexing and centrifugation at 4 °C for 10 min. After centrifugation, the supernatant was transferred to LC-grade vials (ThermoFisher Scientific) and stored at -80 °C until analysis.

### LC-HRMS

In Exp III, to obtain lumen samples for LC-HRMS, the dissected guts (20 guts/sample, four samples) were transferred to microplate wells containing 50 µl of ice-cold M7/well, and the plate was kept on ice thereafter. The guts were allowed to leak out their contents into the ice-cold M7 by applying gentle stirring. After 1-2 min, 50 µl of each lumen content suspended in M7 were transferred to an Eppendorf tube containing 150 µl of ice-cold solvent mixture. The lumen samples were then sonicated for 15 min and centrifuged at 14000 rpm for 10 min at 4°C. The supernatant was then transferred into 320 µL-insert LC -grade glass-vials and stored at -80°C until analysis.

The LC-HRMS analysis were conducted to identify the total (Exp III) and bacteria-produced (Exp IV) L-Dopa. All analytical standards were purchased from Sigma-Aldrich and were of Analytical grades (98-99% purity). Methanol (MeOH), acetonitrile (ACN), and FA were also purchased from Sigma-Aldrich and were of the highest purity (98-99%). MilliQ water was produced in-house using a Milli-Q Integral 3 (LC-Pak polisher, Millipak express 40 filters, 0.22 µm, Merck) for a final organic content < 3ppb. Native standards of L-Dopa and Dopamine were prepared by dissolving the respective powders in MeOH and MilliQ mixtures (80:20), with the addition of 1% (vol/vol) FA, which was then diluted with pure methanol to obtain a 0.1% FA in >98% MeOH solution. These solutions were subsequently stepwise diluted with a 0.1% FA ACN solution to reach the final concentrations of 1.29 and 0.26 µM for the two native standards used for spiking and identification. The standard was prepared in a 1.5 ml glass vial by mixing 800 µl of the standard solution with 200 µl MilliQ and 200 µl ACN.

Instrumental analysis was carried out through injection on the HPLC-HRMS system (Ultimate 3000, ThermoFisher Scientific; Q Exactive HF Orbitrap, ThermoFisher Scientific) with an adapted version of the electrospray ionization settings used in Ribbenstedt et al. [24] (positive ionization, 3700V, sheath gas 30, aux gas 10, sweep gas 0, S-lens RF 50, cap. & aux. gas heater temp 350°C) with data-independent MS2 acquisition (Full-scan: 120k res, max IT 100 ms, AGC target 3e6, scan range 70-1000 m/z; ddMS2: 30k res, max IT 100 ms, AGC target 1e5, loop count 3, TopN 3, isolation window 0.4 m/z, (N)CE 30; ddSettings: Min AGC target 1.00e1, Apex trigger 1-5s, Charge excl. 5-8 & >8, Excl.isot. “On”, Dyn.excl. 2.0s, If idle “Do not pick others”). The system was equipped with an in-line filter (0.5 µm) before the pre-column-fitted HILIC column (Both BEH Amide, 1.7 µm, 2.1 x 5 mm & 2.1 x 150 mm, Waters, USA). The exact mass of L-Dopa was also added to the inclusion list to guarantee MS2 acquisition.

To rule out matrix effects on RT of L-Dopa in the sample matrices and to evaluate intensity drift over the injection sequence, quality Control (QC) samples were prepared by fortifying several already injected replicates of each sample matrix (i. e., lumen and gut microbiome grown on L-Tyrosine) through the addition of 10 µl of three of the calibration mixtures into one replicate each. This way all RT deviations were accounted for, and sequence intensity drift was shown to be 8% over the injections.

All chromatograms were integrated and quantified in XCalibur 3.063 (ThermoFisher Scientific). MS2 spectra were extracted to mzML-format using MSConverter [24] (Peak pick settings: Prefer vendor “check”, MS lvls “1-”; Subset settings: Scan number ““, Scan time ““, Mz win. 0.0-198.10) and compared with the R-script MSMSsim [25] with identities being considered confirmed at a confidence level 1, according to the Schymanski scale [26], when similarity scores > 0.9 and when retention times (RT) in spiked samples matched the samples.

### 16S rRNA Gene Sequencing and Bioinformatics

In Exp IV, the bacterial communities grown at different L-Tyrosine concentrations, including controls, were used for NGS analysis by sequencing the 16S rRNA gene. Genomic DNA was purified with AMPure XP beads (Beckman Coulter) following the manufacturer’s instructions. After the purification, the DNA concentrations were measured using Quant-iT PicoGreen dsDNA Assay kit (ThermoFisher Scientific). Absorbance was measured at 530 nm, using a Tecan Ultra 384 SpectroFluorometer (PerkinElmer). Quality control was performed on an Agilent 2100 BioAnalyser using high sensitivity DNA chip. Libraries were denatured using 0.2 N NaOH and sequenced at LC Sciences (www.lcsciences.com) using the MiSeq Illumina system (2 × 300 bp paired-end) with the v3 reagent kit (600-cycles), following the manufacturer’s instructions. Illumina software v. 2.6.2.1 was used for demultiplexing and removal of indexes and primers according to the standard Illumina protocol. Following demultiplexing, removal of tags and primers, the reads were processed using the *DADA2* package (version 1.6) as implemented in the R statistical software v. 3.4.2 (R Core Team, 2016) that detects and removes low-quality sequences and merges paired-end reads to generate amplicon sequence variants (ASVs). The sequences were trimmed 60 bp downstream of the forward primer and 100 bp downstream of the reverse primer. Sequence quality control was conducted by identifying and removing chimeric sequences and those containing ambiguous bases. The resulting sequences were classified with the SILVA taxonomy (Silva Ribosomal RNA database; version v.132). Sequences have been deposited with links to BioProject accession number PRJNA694094.

Illumina MiSeq sequencing of all amplicons resulted in a total of 647 891 high-quality filtered reads in the bacterial samples, with a mean read depth per sample of 43 192 sequences and a total ASV number of 120 (49 with ≥ 2 counts). Data filtering was applied using a minimum count of 4 and 20% prevalence to remove low-quality or uninformative features. Due to the moderate variability in the sequence libraries, the data were not rarefied for diversity analysis. Rarefaction curves and Zhang-Huang’s coverage estimator (Fig 3c) were calculated from ASV abundances using functions supplied by the *vegan* and *entropart* R packages.

### Data Analysis

#### Evaluation of the differences in the *Ddc* gene expression among the molt stages

As the gene expression data were collected using different cohorts on three occasions, we normalized data by Z score transformation, using mean value and standard deviation for each cohort, to standardize data across the experimental runs. The normalized data were independent of the absolute variation between the individuals from different cohorts and used to compare groups (postmolt, intermolt, and premolt) by the pairwise multiple comparison Holm-Šídák test with multiplicity-adjusted *p* values [27]. The null hypothesis was rejected with a probability of error α < 0.05.

#### Microbial community structure and identification of taxa responding to L-Tyrosine

To visualize the differences in the bacterial community structure, a heat-map with cluster analysis was used at the genus level (> 0.2%). The R-package *edgeR* was used to identify differentially abundant bacterial taxa based on the false discovery rate-corrected *p*-values (α = 0.05, FDR = 1%) that were associated with controls and L-Tyrosine incubations. PHYML in Geneious Prime® 2019.2.3 was used to reconstruct phylogenetic tree using 16S rRNA gene sequences of all treatments. The bacterial communities obtained in the L-Tyrosine incubations (Fig. S3) were representative of the *D. magna* gut microbiota for this clone [11]. All figures were prepared using BioRender (https://biorender.com).

## Results

### Localization of peripheral CA pathway in the gut

TH and DDC enzymes were localized in the daphnid midgut wholemount (Fig 2a) using immunohistochemistry (Exp I), suggesting a peripheral CA-pathway in the gut wall. TH presence (Fig. 2b-c) indicates hydroxylation of L-Tyrosine to L-Dopa, which is the first and critical limiting step of CA-pathway, whereas DDC presence (Fig. 2d-e) indicates decarboxylation of L-Dopa to Dopamine, the second step (Fig. 1). The occurrence of TH and DDC in the gut wall, including the microvilli layer, suggests a local Dopamine synthesis associated with the gut wall, including epithelium.

**Figure 2.**
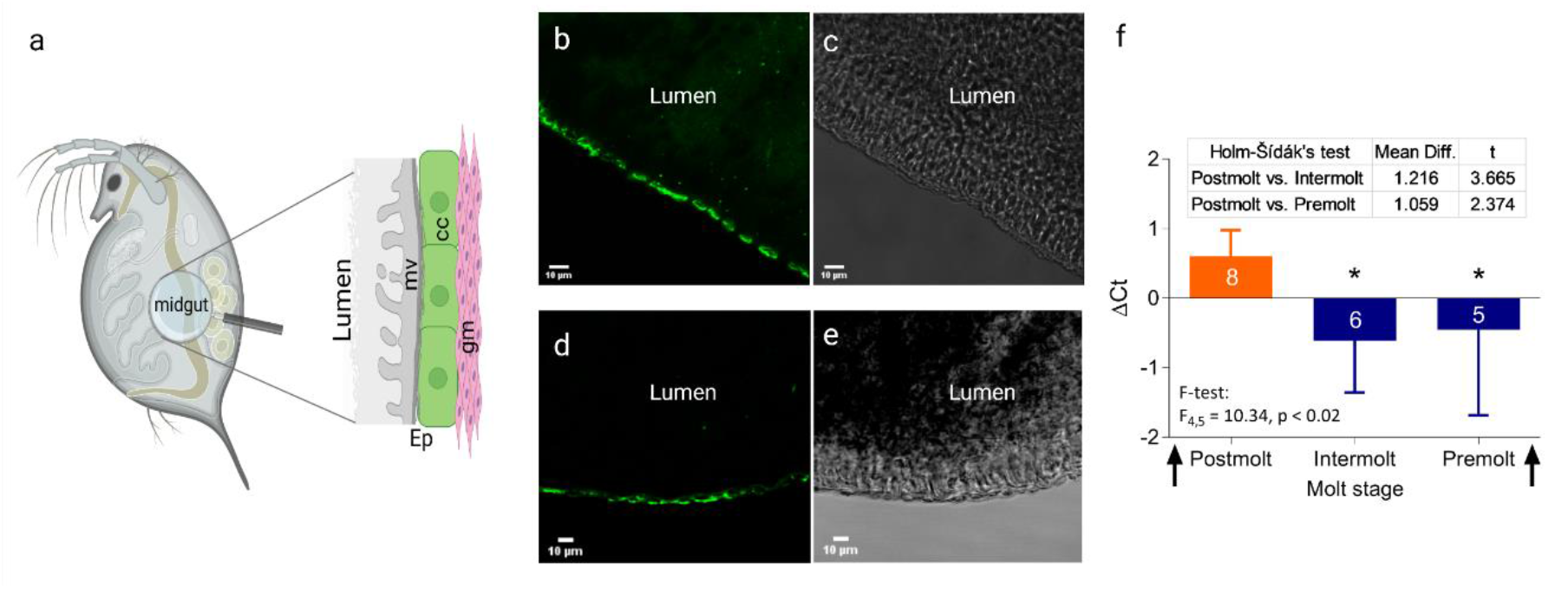
L-Dopa production in *Daphnia magna* gut. **a)** Schematic structure of the midgut [49] analyzed with immunostaining (*mv* – microvilli layer; *cc* – columnar cells; *gm* – gut musculature; *Ep* – gut epithelium comprised of *mv* and *cc*); **b&c**) Immunofluorescence localization of TH in the mid-gut epithelial cells; **d&e)** Immunofluorescence localization of DDC in the mid-gut epithelial cells; the same region as for TH but in a different individual. Labeling with Alexa Fluorophore 488 is shown on the left and the light microscopy of the same specimen on the right; no labeling appeared in the unlabeled control (not shown); **f)** The significant upregulation of *Ddc* gene expression in *Daphnia* gut observed during the postmolt decreasing during the intermolt and premolt; arrows indicate ecdysis. The data are shown as z scores (mean ± SD, number of observations are shown on the bars for each stage) normalized to the cycle-specific average values for each of the replicate experimental run.

### Dynamics of *Ddc* expression during the molt cycle

In Exp II, we used qPCR assay to measure the *Ddc* gene expression in the daphnid gut during the molt cycle with ecdysis as a reference point (i.e., postmolt, intermolt, and premolt). The *Ddc* gene was expressed in all samples tested (Fig. 2f), indicating that the DDC synthesis is continuous. Moreover, there was a significant association between the *Ddc* expression and the daphnid molt stage, with the highest values observed in the postmolt animals (pairwise comparison with the Holm-Šídák test; F4,5 = 10.34, *p* < 0.02; Fig. 2f).

### L-Dopa in the lumen

In Exp III, we used liquid chromatography high-resolution mass spectrometry (LC-HRMS) to identify CA in *Daphnia* lumen samples prepared from the dissected guts. Peaks of the exact mass corresponding to L-Dopa (<3 dPPM) were detected in all samples, and spectra fragmentation revealed a similarity of 0.905 with a native L-Dopa standard (Fig 3a-b), allowing positive L-Dopa identification [26]. Dopamine was not detected in these samples at LOD of 0.47 pg/µL; therefore, it was either below the detection limit or conjugated and biologically inactive [28].

**Figure 3.**
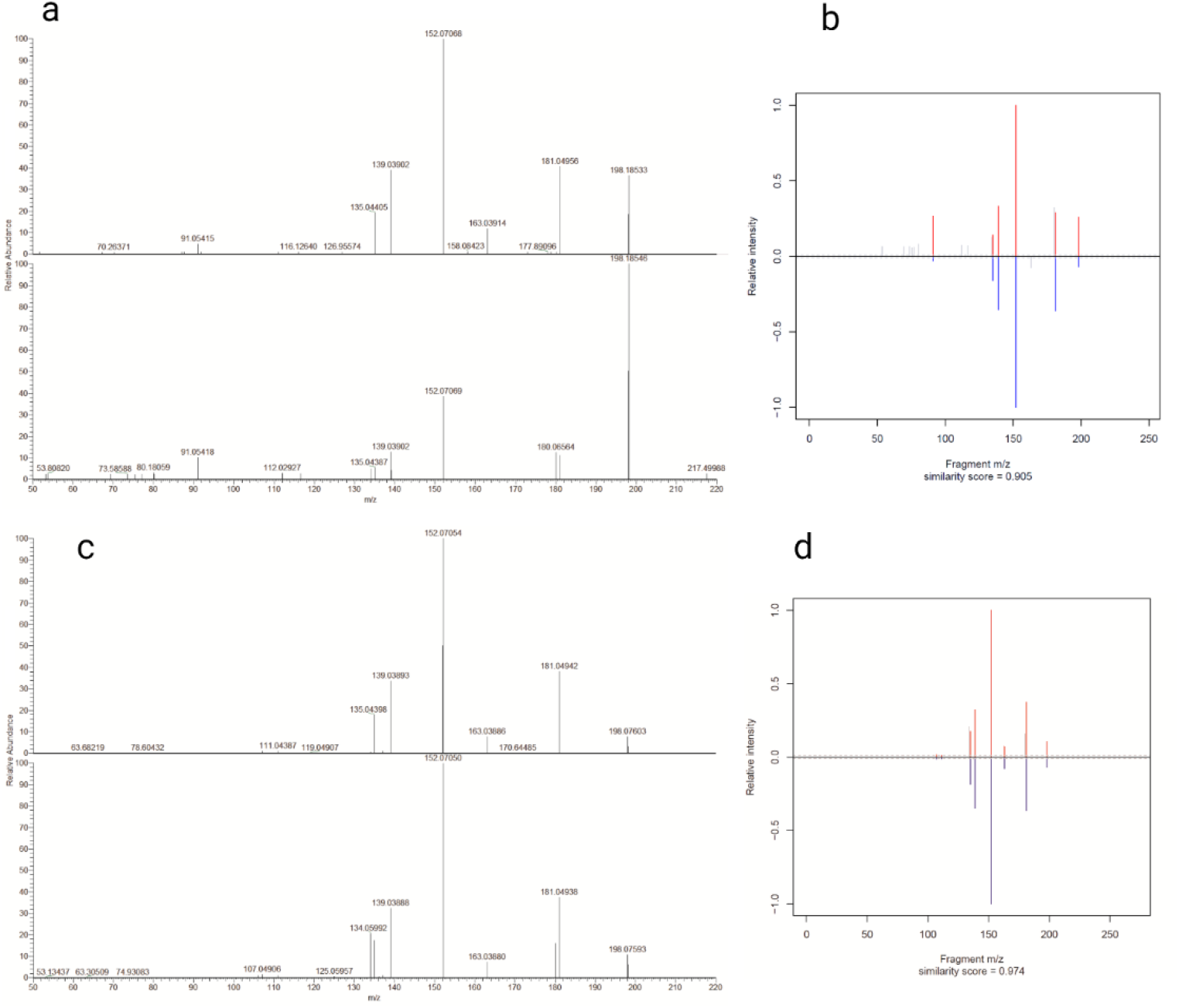
Presence of L-dopa in lumen and its production by the enriched gut microbiome of *D. magna*. **a)** Identification of L-Dopa in *Daphnia* lumen by comparing fragmentation pattern of the native standard (upper spectrum) to the endogenous L-Dopa in the lumen (lower spectrum); **b)** MSMSsim similarity score for peaks with exact mass of L-Dopa from the *Daphnia* lumen *vs*. the standard; **c)** Identification of L-Dopa produced by the microbiome *in-vitro* (2 mM L-Tyrosine treatment) by comparing fragmentation pattern of the native L-Dopa standard (upper graph) and bacteria-produced L-Dopa (lower graph); **d)** MSMSsim similarity score for peaks with the exact mass of the bacteria-produced L-Dopa and the standard.

### Production of L-Dopa by gut bacteria

The L-Dopa detected in the lumen could be produced by the host (as shown in Exp I) and/or the microbiota that utilizes foodborne L-Tyrosine for L-Dopa synthesis [7]. To evaluate whether the daphnid microbiota is capable of L-Dopa production, we conducted an enrichment experiment *in-vitro* by amending gut bacteria with L-Tyrosine at 0 to 3 mM concentration range (Exp IV). Bacterial growth responded to the enrichment in a dose-dependent manner (based on OD600 dynamics; Fig. S1). Spectral fragmentation of the L-Dopa produced by the bacteria revealed spectral similarities above 0.949 with native L-Dopa standard at ≥2 mM L-Tyrosine (Fig. 3c-d). Therefore, L-Dopa detected in the *Daphnia* gut lumen can originate from both the host and its microbiota.

### Bacterial taxa associated with L-Dopa production

We conducted NGS analysis to identify bacterial taxa associated with the L-Dopa production in the L-Tyrosine amended gut microbiota. The 16S rRNA gene libraries were obtained by Illumina MiSeq sequencing using the gut microbiota obtained in the L-Tyrosine enrichment experiment (Exp IV). The differential abundance analysis showed that *Pseudomonas, Akkermansia*, and *Butyrivibrio* were upregulated in the L-Tyrosine exposure, with significant upregulation in *Pseudomonas* (Fig. 4).

**Figure 4.**
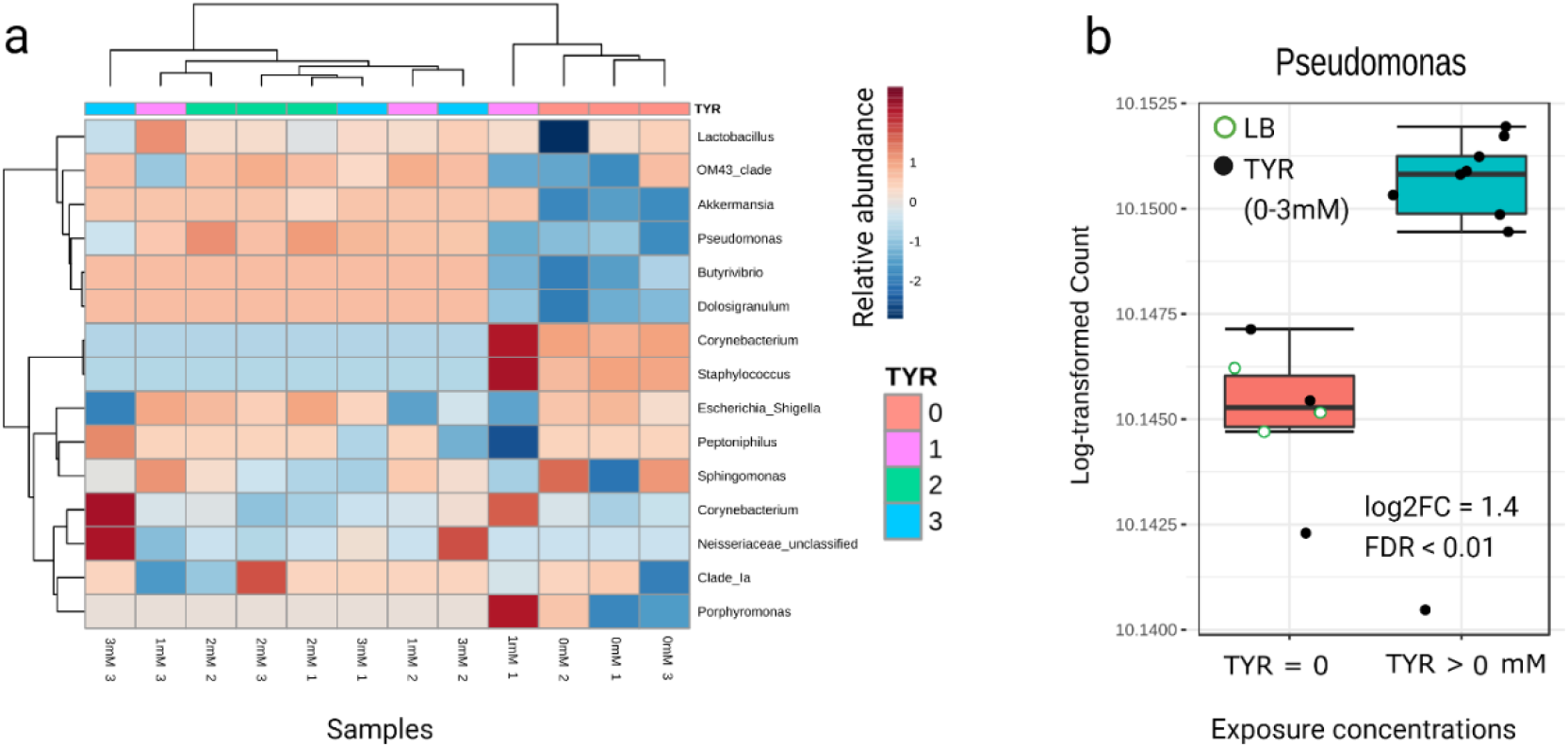
Identification of taxa producing L-Dopa. **a)** Heat-map with cluster analysis at the genus level (>0.2%) for relative abundance; and **b)** results of the differential abundance analysis for significantly upregulated taxa in the L-Tyrosine treatments showing that *Pseudomonas* was significantly associated with the L-Tyrosine exposure.The samples were grouped to show abundances from 0 and LB (TYR = 0) and 1 to 3 mM (TYR > 0 mM) treatments. See Fig S3 for the composition of the bacterial community in the experiment.

## Discussion

We demonstrated for the first time that the neurotransmitter L-Dopa is produced in the gut jointly by *Daphnia* and its microbiome and used for Dopamine synthesis in concert with the molt cycle progression of the host. Moreover, immunostaining of TH and DDC in the gut wall suggests a peripheral CA pathway in *D. magna*. To the best of our knowledge, no cells in the gastrointestinal tract that contain Dopamine and express components of Dopamine-signalling pathways, including enzymes and specific Dopamine receptors and transporters, have been reported in *Daphnia*. In decapod crustaceans and insects [29–32], dopamine and other neurotransmitters are present in the gut innervation, contributing to gut motility regulation, but there are only a few studies on the neuroendocrine cells associated with gut lining and their functioning. Notably, in *Drosophila* larvae, only 6% of DDC activity is associated with the brain; the rest occurs in the epidermis [33]. Moreover, enteroendocrine cells account for 5-10% of the midgut epithelial cells in flies [34]. In the gastrointestinal tract and other mesenteric organs of vertebrates, Dopamine production is substantial [35, 36], and gut epithelial cells contain DDC and receptors for L-Dopa with both endogenous L-Dopa and lumen L-Dopa involved in Dopamine synthesis [37].

The central nervous system in the water fleas and other lower crustaceans consists of the brain, compound eye, optic ganglia, and thoracic nervous system [18, 19, 38]. To visualize neuropils, neurosecretory somata, and neurotransmitter-producing neurons, including histaminergic [18], peptidergic [39], and dopaminergic neurons [17], immunostaining is commonly used. However, we know much less about the peripheral nervous system in these model species, especially the gut innervation, and no sensory neurons in the gut have been visualized in *Daphnia* [18]. Histaminergic somata have been identified along the neuropils extended from the dorsal and ventral cords surrounding the gut, but none were seen in the periphery reaching the gut wall [18] as shown for *Artemia* [40]. Our immunohistochemical analysis (Fig 2 a-e) has revealed relatively homogenous labeling pattern along the gut wall. Given the observed staining of microvilli layer, we suggest that TH and DDC detected in our study are produced by the gut lining. However, this does not exclude the possibility that both neurotransmitter-producing somata and dopaminergic fibers similar to the serotoninergic fibers associated with the gut musculature [41] in insects are involved.

The release mechanisms for L-Dopa and Dopamine in the cuticle of *Daphnia* or other crustaceans and their ecological triggers are poorly understood. We found free unconjugated and hence biologically active L-Dopa in the *Daphnia* lumen, but no Dopamine, which supports the suggested pathway (Fig. 1). However, the differences in the transport systems between L-Dopa and Dopamine may also contribute. Both molecules are transported using different systems adapted to their chemical structure. In neurons, Dopamine is transported via the specialized vesicular monoamine transporter (VMAT) or Dopamine transporter (DAT) [42, 43], while L-Dopa is transported via the large neutral amino acid transporter (LNAA) system [44], a heterodimeric membrane transport protein that preferentially transports branched-chain and aromatic amino acids. Thus, a lack of a specialized transport system may explain why we detected L-Dopa but not Dopamine in the lumen.

The *Ddc* gene is evolutionary highly conserved across taxa [45, 46]. We followed the expression of *Ddc* in the gut and found that it follows the molt cycle, peaking after ecdysis. DDC is an enzyme found in metazoans [45], whereas bacteria use other pathways to transform L-Dopa to Dopamine [47]; hence, the expression pattern reflects the daphnid *Ddc* expression. According to qPCR results and confocal microscopy, transcription and biosynthesis of DDC occur in the gut wall, which is in line with a correlation between epidermal *Ddc* expression and the protein activity reported in insects, with no translational modification [45]. Dopamine and its derivatives are essential for arthropod physiology, including gut motility, cuticle formation, and sclerotization of the integument during the postmolt [41, 46], and thus involved in the basic functions, i. e. feeding, ontogenetic development, behavior, wound-healing, and protection against pathogens [45]. As feeding and cuticle formation are crucial for growth and development and tightly regulated by the molting hormone ecdysone, *Ddc* is also under tissue-specific and hormonal control [45, 48]. In line with our *Ddc* expression results, epidermal *Ddc* activity in *Drosophila* peak prior to embryo hatching, during larval-larval molting, pupariation, and adult eclosion [45, 48]. For cuticle sclerotization, DDC is a rate-limiting enzyme, and mutation loss of *Ddc* in *Drosophila* is lethal at the embryonic stage [42]. Therefore, the cuticle formation in *Daphnia* foregut and hindgut [49] is a possible target for the *Ddc* gene expression and the associated DDC production peaking shortly after ecdysis. However, other targets and processes following the molt cycle can be involved.

Besides facilitating digestion, gut microflora participates in host metabolism and behavior through their ability to produce, recognize and modulate eukaryotic neurotransmitters and other info-signals [2]. We found that the gut microbiota of *Daphnia* is capable of L-Dopa production *in vitro* following enrichment with L-Tyrosine. Moreover, the endogenous *Pseudomonas* spp. was significantly upregulated in a concentration-dependent manner. In line with this, several bacterial taxa have been reported to produce L-Dopa, including *Pseudomonas* spp. [50], and several bacterial enzymes for L-Dopa production have been identified [7]. We suggest that *Daphnia* and its gut bacteria are engaged in host-microbiome communication using L-Dopa as a signaling molecule. Our findings that L-Dopa is produced by both the gut epithelium and the microbiota and reports showing that bacteria recognize and respond to L-Dopa [6] support this hypothesis. In turn, as a grazer, *Daphnia* can use chemosensation to modulate feeding behavior and food uptake in response to amino acids present in the feeding environment [51]. Tyrosine and its precursor phenylalanine were found to stimulate *D. magna* mandible movements by 43% and 34%, respectively [51], which would provide an increased supply of this amino acid to the microbiota and the substrate for L-Dopa synthesis.

In summary, TH and DDC are present in the *Daphnia* gut wall, and both the host and its microbiota contribute to the free unconjugated L-Dopa in the gut (Fig. 1). Therefore, L-Dopa is a putative agent for host-microbiome communication in daphnids and, perhaps, other invertebrates translating into the ecologically important host traits. In invertebrates, gut bacteria affect epithelium development [42, 43], modulate growth factor signaling, and gut stem cell activity [52]. The fact that *Ddc* is always expressed, peaking after ecdysis, suggests continuous Dopamine synthesis, its possible involvement in active feeding during the postmolt, cuticle formation, and animal development. By contributing to L-Dopa production, the gut bacteria may affect Dopamine synthesis and host behavior. In turn, by modulating the amino acids and, particularly, Tyrosine intake, the host might benefit by regulating microbial L-Dopa synthesis.

Our work provides new insights into the molecular mechanism(s) by which lower crustaceans communicate and interact with their microbiome and increase our fundamental knowledge about the role of L-Dopa, not as a Dopamine precursor as conventionally assumed, but as an agent of inter-kingdom communication in crustacean ecophysiology and regulation of ecological traits. Further work will identify the biochemical, physiological, and ecological context for the host-microbiome interactions conveyed by lumen L-Dopa and explore its commonality across phylogenies.

## Acknowledgments

This research was supported by The Swedish Research Council (VR) for BioDeg project [grant number 2018-05213] and The Swedish Research council for Sustainable Development (FORMAS) [grant number 2018-01010] to EG and RE.

## Author Contributions

Middle authors contributed equally. RE developed the hypothesis with contribution from EG, participated in NGS bioinformatics analysis, and wrote the first draft. Corresponding author EG contributed to writing and performed bioinformatic and statistical analyses with some contributions from RE. Authors RE, SL-J, and AR conducted the laboratory experiments. Author AR conducted LC-MS/MS data analysis.

